# NOK promotes tumorigenesis through coordinating epidermal growth factor receptor to boost the downstream signaling in breast cancer

**DOI:** 10.1101/2024.08.14.608018

**Authors:** Yinyin Wang, Bingdong Zhang, Chunhua He, Bo Tian, Sihan Liu, Jianghua Li, Jiayu Wang, Shigao Yang, Bingtao Zhu, Xiaoguang Wang, Zhijie Chang, Chenxi Cao

**Affiliations:** State Key Laboratory of Membrane Biology, School of Medicine, Tsinghua University, Beijing (100084), China; Department of Gastrointestinal Surgery/Department of Clinical Nutrition, Beijing Shijitan Hospital, Capital Medical University, Beijing (100700), China; Department of Surgery, The Second Affiliated Hospital of Jiaxing University, No. 397, Huangcheng North Road, Jiaxing, Zhejiang (314000), China; Department of Surgical, Hospital of Northwestern Polytechnical University, Xian, (710072), Shaanxi, China; School of Life Sciences, Anhui Medical University, Hefei 230032, China

**Keywords:** EGFR signaling, NOK/STYK1, STAT3, STAT5, breast cancer

## Abstract

**BACKGROUND:** Epidermal growth factor receptor (EGFR) forms a homodimer or heterodimer with other ErbB receptor family members to activate different downstream cytoplasmic signaling proteins during tumorigenesis.

**METHODS:** Adenovirus and lentivirus were used to overexpress or deplete NOK and/or EGFR to evaluate the phosphorylation of EGFR, the interaction of NOK-EGFR and their role in cell proliferation and metastasis.

**RESULTS:** EGFR heterodimerizes with NOK (also known as STYK1), a novel tyrosine kinase with a transmembrane domain, to promote tumorigenesis and metastasis of breast cancer cells. We found that NOK directly interacted with EGFR and formed a heterodimer complex. Depletion of NOK impaired, but over-expression of NOK increased, the phosphorylation of EGFR. NOK enhanced EGF signaling activation, in particular, the phosphorylation of STAT3, STAT5 and Erk1/2 via its juxtamembrane (JM) domain in promoting the proliferation and migration of breast cancer cells. Overexpression of NOK and EGFR synergistically induced the tumorigenesis of NIH-3T3 normal cells. We finally demonstrated that co-expression of NOK and EGFR correlated with tumor malignant stages in breast cancer patients.

**CONCLUSIONS:** Our findings uncover a mechanism by which NOK coordinates EGFR to enhance EGF signaling during tumorigenesis and metastasis and propose a potential strategy for targeting NOK-EGFR in breast cancer treatment.

**GRAPHICAL ABSTRACT:** 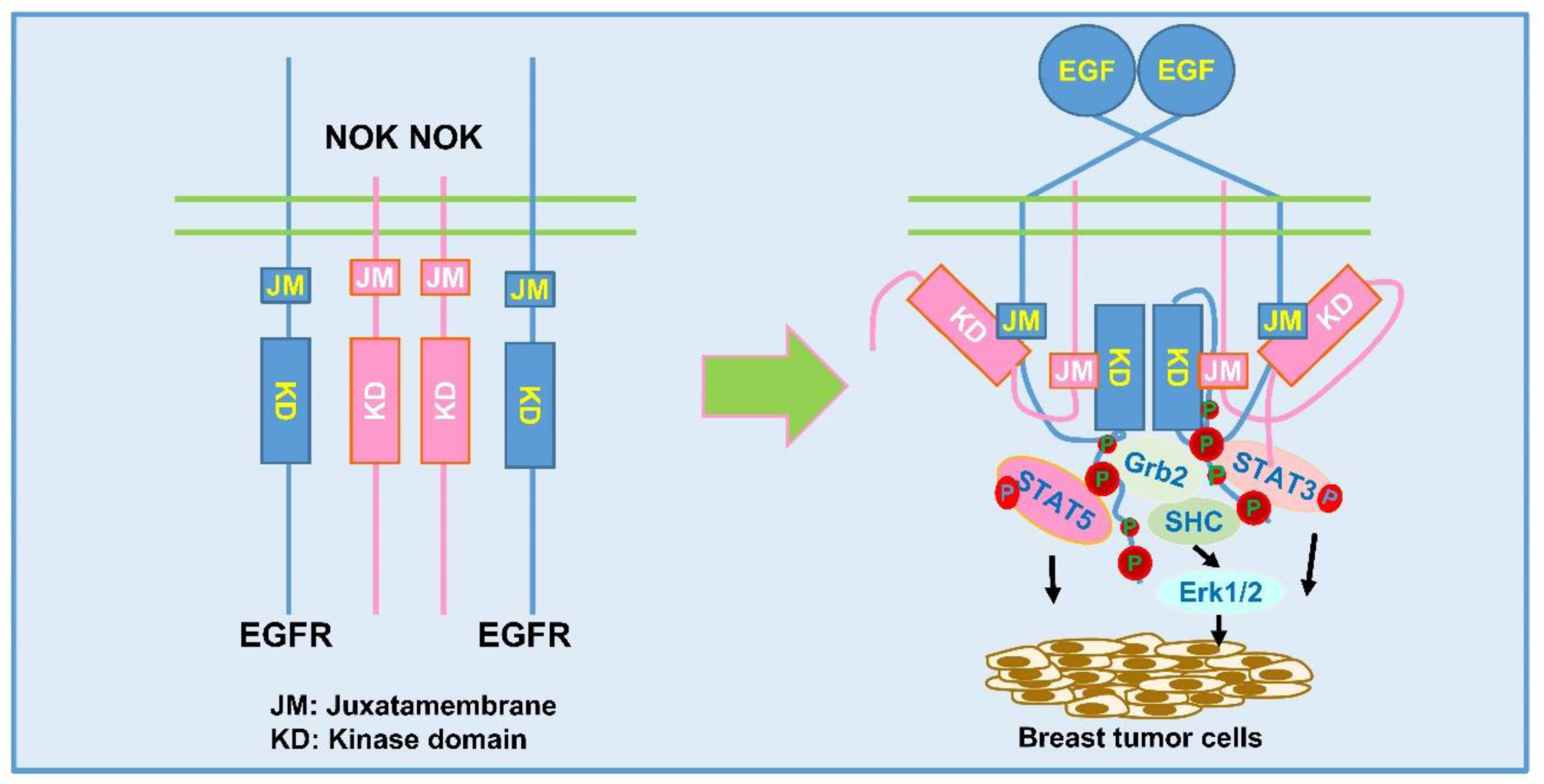

1. EGFR heterodimerizes with NOK/STYK1, a novel tyrosine kinase with a transmembrane domain, in a manner of cross interaction via their juxtamembrane (JM) domains and kinase domains.
2. NOK enhances EGF signaling activation, in particular, the phosphorylation of STAT3, STAT5 and Erk1/2 via its JM domain.
3. NOK and EGFR synergistically promote proliferation and migration of breast cancer cells and induce tumorigenesis of normal cells.
4. Co-expression of NOK and EGFR correlates with tumor malignant stages in breast cancer patients.

## BACKGROUND

EGFR, a member of the ErbB family of receptor tyrosine kinases (RTK), plays an important role in the regulation of cell proliferation and migration [1]. EGFR owns an extracellular domain (ECD), a hydrophobic transmembrane domain (TM), an intracellular catalytic tyrosine kinase domain (KD) and several intracellular tyrosine residues [2, 3]. The epidermal growth factor (EGF) family of peptide growth factors activate EGFR by binding to its extracellular domain and inducing formation of a dimer complex. Dimerization of EGFR leads to auto-phosphorylation of the tyrosine residues in the intracellular domain of one EGFR monomer by the catalytic site of the other monomer [4]. The phosphorylation sites then recruit adapter proteins and other signaling molecules for signal transduction. Downstream the receptor, several effectors, including Ras, MAPK, Src, STAT3, STAT5, PLC-3, PKC and PI3 kinase, are activated to relay signal from the cytoplasm into the nucleus [1]. Importantly, the specific activation of the downstream factors depends on the status of EGFR dimerization and the subsequent recruitment of effectors [5, 6].

EGFR cross-talks with other receptors. EGFR interacts with PDGFR, IGF-1R, Axl RTKs and c-MET [7–11]. The interaction of EGFR and PDGFR enables PDGF-induced trans-activation of the EGFR signaling in aortic vascular smooth muscle cells [10]. EGFR and IGF1R physically interact to protect IGF1R from ubiquitination and proteasome degradation. Yet, the association of EGFR and IGF1R fails to trans-activate each other in breast cancer cells [9]. EGFR also interacts with Axl physically at the cell membrane and directly phosphorylates Axl [8]. Another study indicated that EGFR was co-expressed with c-Met in SUM229 cells, where EGFR coordinated c-Met to recruit c-Src, maintaining the constitutive phosphorylation of c-Met [12]. All these studies indicated that the association of EGFR with other receptors is also critical for tumor proliferation. However, it remains interesting to identify more interacting receptors for differential regulation of EGFR.

NOK, also known as a putative serine/threonine and tyrosine receptor protein kinase (STYK1), was identified by our group as a member of the RTK family [13]. Our previous studies showed that NOK induced transformation, tumorigenesis and metastasis [13, 14]. To date, NOK has been reported to be abundantly expressed in breast (18), ovarian (19), colon (20), prostate, lung cancers [15, 16] and acute leukemia (22). The elevated expression of NOK was attributed to its regulation of m(6)A modification by RBM15 [17] and the transcription by Smad3 [18]. These studies indicated that NOK could be a potential marker for cancer diagnosis (18, 23). Functionally, others reported that overexpression of NOK increased cell proliferation in castration-resistant prostate cancer, as well as epithelial-mesenchymal transition and tumor metastasis in human hepatocellular carcinoma [19]. Reciprocally, depletion of NOK was reported to inhibit cell proliferation, tumor formation and invasion in intrahepatic cholangiocarcinoma and glioma cells [20, 21]. Mechanistic studies demonstrated that NOK promoted metastasis and epithelial-mesenchymal transition by inhibiting FoxO1 signaling [22] and regulated ferroptosis [23] in non-small cell lung carcinoma. Recent studies reported that NOK functioned in immune cells and regulated tumor metabolism and microenvironment [24–26]. This echoes its role on the regulation of TGF-β1 in the laryngeal squamous cell carcinoma [27]. Importantly, as a transmembrane protein, previous studies demonstrated that NOK activated several signal pathways including RAS/MAPK, PI3K/Akt and STAT1/STAT3 [19, 28–31]. However, NOK displayed a very weak kinase activity when ectopically expressed in cells [13], although it was recently reported that EGFR mediated phosphorylation of NOK [32]. Nevertheless, several studies indicated that NOK functioned together with EGFR in lung cancers [32–34]. These studies implied that NOK might participate the signal transduction of EGFR. However, how NOK assists EGFR phosphorylation remains unclear. In this study, we report that NOK interacts with EGFR via the JM domain of NOK and kinase domain of EGFR to activate STAT3 and STAT5 to synergistically promote tumorigenesis of breast cancer cells..

## METHODS

### Cell culture and stable cell line construction

MDA-MB-231 and HEK293T cells were obtained originally from the American Type Culture Collection. All media were from Invitrogen and supplemented with 10% fetal bovine serum (Invitrogen, Pleasanton, CA, USA) and cells were maintained at 37°C in an atmosphere containing 5% CO2. MEF cells were acquired from the day 13 embryo of wildtype or NOK knock out mice (sent as a gift by Dr. Dong Zhongjun, Tsinghua University, China, data unpublished) and used in four generations. Lentivirus was used to establish stable cell lines for over-expression and depletion of EGFR, NOK, and the mutants in this study. Cells were transiently transfected with the indicated plasmids using Lipofectmine2000 (Invitrogen, CA, USA) according to the manufacturer’s instructions. Mycoplasma contamination was monitored using MycoFluor mycoplasma detection kit (M7006, Invitrogen, CA, USA).

### Plasmids

EGFP-N3/NOK, EGFP-N3/NOK-ICD(49-422), and EGFP-N3/NOK-KD(106-381) were constructed in our lab [35]. EGFP tagged NOK mutants were constructed by using EGFP-N3/NOK as a template, with HindIII/SalI as cloning sites. Flag-EGFR was a gift from Dr. Lan Ma (Shanghai Jiaotong University, China). EGFP-N3/EGFR was kept in our lab [36]. Different mutants of EGFR were constructed using EGFP-N3/EGFR, with HindIII/XhoI as cloning sites. EYFP-N1/EGFR was constructed from EGFP-N3/EGFR. ECFP-N1/NOK was constructed using EGFP-N3/NOK. Ad-NOK was constructed following a previous report [37] with BglII/XhoI as cloning sites. pCDH-3HA/NOK and all the pCDH-3HA/NOK deletions were constructed using GFP-N3/NOK. ELK-Luc, APRE-luc, and LHRE-luc reporters were kept in our lab [38]. Ad-track/NOK was constructed according to a previous protocol[37]. Ad-easy vector and Ad-track/GFP were maintained in the lab.

### Antibodies and reagents

Antibodies against FLAG (M2, F3165), β-Actin (A2228), and NOK (SAB1302051), and all other chemicals were purchased from Sigma (St. Louis, MO, USA). Antibodies against Myc (9E10, sc-40), GFP (FL, sc-9996), HA and HA-HRP (F-7, sc-7392), STAT5B (sc-1656), phospho-ERK1/2 (sc-81492) and Grb2 (sc-8034) were purchased from Santa Cruz Biotechnology (Dallas, TX, USA). Antibodies against phospho-EGFR (Tyr-1173, 4407), phospho-EGFR (Tyr-845, 6963), phospho-STAT3 (Tyr-705, 9131), STAT3 (9139), ERK1/2 (9102), EGFR (4267), phospho-STAT5 (Tyr694/Tyr699, 9359), phospho-AKT (Ser-473, 9271), AKT1/2/3 (9272) and SHC (2432) were purchased from Cell Signaling (Danvers, MA, USA). Antibody against phosphotyrosine (4G10, 05-321) was purchased from Millipore (Billerica, MA, USA). EGF (324831, Calbio-chem, Germany) was purchased from Bio-Chem Industries (Sapulpa, OK, USA).

### Depletion of NOK using RNA interference

NOK was depleted with either shRNAs or siRNS. shRNAs for depletion of NOK and control shRNA for scramble (NC) [39] were synthesized with two single-strand fragments, inserted into XbaI-XhoI sites of the pLL3.7 vector for lentivirus constructs, which were used for stable cell selection as described previously [40]. For luciferase assay or Western blot, two siRNA sequences targeting different regions of NOK were designed as #1: 5’-GCAAGAAACAUUCAUGCAU-3’ and #2: 5’-GUCUUUCCCAGGGACACAA-3’ (SASI_Hs01_00043410 and SASI_Hs02_00351937, Sigma, St. Louis, MO, USA). The control siRNA is a scrambled sequence (5’-UUCUCCGAACGUGUCACGUdTdT-3’) (NC, Sigma, St. Louis, MO, USA). A total of 60 nM siRNAs was used for transfection using Lipofectamine-RNAi-MAX Reagent (13778, Invitrogen, CA, USA) according to the manufacturer’s instructions.

### Luciferase reporter assays

Indicated plasmids were co-transfected into the cells with TK as an internal control. After transfection for 24 h, cells were starved with DMEM containing 0.2% FBS for 12 h and then stimulated with EGF (50 ng/ml) for 8 h. The reporter activity was examined by the Dual-Luciferase Assay System (T002, Vigorous Inc., Beijing, China). Firefly luciferase activity was normalized against Renilla luciferase activity and presented as a mean ± standard deviation (SD).

### Immunoprecipitation (IP) and Western blot

Cells were subject to starvation for 12 h and treated with 100 ng/ml EGF for 10 min or indicted times. The cells were lyased by RIPA lysis buffer for 30 min. Western blot was performed according to a previous protocol [41]. For the IP assay, the cell lysis was incubated with the indicated antibodies overnight at 4℃.

### Cell proliferation assay

Different cells were seeded into 96-well plates in a density of 1000-1200 cells/well. After cell attachment, low serum medium (2% serum) was changed overnight and then EGF (50 ng/ml) was added for the indicated times. The number of cells was counted at different times using Cell Titer 96 A Queous Non-radioactive Cell Proliferation Assay kit (G5421, Promega, Madison, WI, USA).

### Adenovirus-driven NOK expression and soft agar colony-forming assay

Ad-NOK and Ad-GFP were packed using HEK293 cells. MDA-MB-231 cells infected with Ad-GFP or Ad-NOK for 24 h (1.0×10^3^ per dish) were re-suspended in 0.35% agar with or without EGF (50 ng/ml) and then seeded on the top of a 1 ml solidified 0.6% agar layer in 35-mm dishes. DMEM with or without EGF (50 ng/ml) was added to the top layer of 0.35% agar. Colonies >80 μm in diameter were counted after 14 days.

### Colony formation assay

Cells were seeded into 6-well plates (1000 cells per well) and cultured with or without EGF (50 ng/ml) for 14 days, then washed with PBS and stained with 0.1% crystal violet. The number of colonies was counted by ImageJ and presented as the mean ±standard deviation (SD) from three individual experiments.

### Wound healing assay and cell invasion assay

Wound healing assay was performed as a previous report [40]. Cells were seeded into 6-well plates and cultured for 24 h. The monolayer cells were wounded with a sterile plastic tip. After washing twice with PBS, the cells were treated with or without EGF (50 ng/ml) for 24 or 36 h. Cell migration was observed by a microscopy and relative migration rates were measured by ImageJ and quantified by a formula as relative distance of migration (%) = (distance of oh-distance of 30 h)/distance of 0 h*100% and presented as means ±S.D.

### Cell invasion assay

The *in vitro* invasive ability was evaluated using a transwell chamber of 8 μm pore size (Millipore Corp., Bedford, MA, USA) coated with 100 ul of Matrigel (BD Biosciences, Franklin Lakes, NJ, USA, 1:10 diluted). Cancer cells (1 × 10^6^) were re-suspended in 300 μl 0.1% serum-free medium and cultured in the upper chamber. The lower chamber was filled with 200 ul 10% FBS medium (with or without 50 ng/ml EGF). The filters were fixed with methanol for 10 min and stained with 0.5% crystal violet (0.02% in PBS) for 30 min. Cells on upper surfaces of the filters were removed with a cotton swab. Cells on the reverse sides were counted in three random fields using a microscope at a 100× magnification. Each assay was performed in triplicate.

### Immunofluorescence analysis and fluorescence resonance energy transfer (FRET) assay

Cells were fixed with 4% paraformaldehyde for immunostaining using the indicated antibodies, followed by incubation with secondary antibodies conjugated with FITC or TRITC (Jackson Research Laboratories, Bar Harbor, ME, USA) [36]. FRET was performed in living cells using the acceptor photo-bleaching method by a confocal laser scanning microscope (A1Rsi; Nikon, Inc., Chiyoda, Japan) [41].

### Xenograft tumor model

MDA-MB-231 cells were infected with Ad-GFP or Ad-NOK (2 ×10^6^ in 100 μl) or stably transfected with shRNAs against NOK (5 × 10^6^ in 100 μl). NIH-3T3 cells were stably transfected with EGFR or infected with Ad-GFP or Ad-NOK (5 × 10^6^ in 100 μl). All the cells were subcutaneously inoculated in the flanks of BALB/c-nu/nu mice (5 weeks). Five or three mice were used in each group. Tumor volumes were monitored every 3 days and calculated using the following formula: volume = (length × width^2^)/2. After about 3-8 weeks mice were sacrificed and tumors were excised and weighed. All the animal experiments were performed with the approval of the Animal Healthy and Ethic Board of Tsinghua University.

### Lung metastasis experiment

MDA-MB-231 cells infected with Ad-GFP or Ad-NOK (5 × 10^6^ in 100 μl) were injected into BALB/c-nu/nu mice via the tail vein. After 60 days, mice were euthanized, and metastatic nodules on lungs were stained with Boulin’s solution and counted.

### Immunohistochemistry

Tissue microarray slides, purchased from Shanghai OutDo Biotech Co. LTD (HBre-Duc170Sur-01, Shanghai, China) were used for immunohistochemistry analyses with a monoclonal antibody against NOK, a polyclonal antibody against EGFR and monoclonal antibodies against pSTAT5, pSTAT3 or Ki67 [42].

### Statistical analysis

All data were representative of at least three independent experiments. Statistical significance of differences between mean values was assessed with paired Student’s t-test. Minimum level of significance was set at 0.05.

## RESULTS

### NOK enhances growth and metastasis of breast cancer cells in nude mice

To address whether NOK is related to breast cancers, we analyzed its expression pattern in tumor and normal tissues using data from The Cancer Genome Atlas (TCGA) and Genotype-Tissue Expression (GTEx) databases (https://commonfund.nih.gov/GTEx/). The result showed that NOK was highly expressed in tumors compared with normal tissues (Fig. 1A) and the expression of NOK was significantly negatively correlated with the survival of the breast cancer patients (Fig. 1B). To verify the result from these databases, we performed a western blot and IHC staining analysis using clinical samples. The result showed that NOK was highly expressed in most of breast cancer tumors compared with the adjacent tissues in 4 patients (Fig. 1C). To examine whether NOK functions on tumor progression and metastasis, we over-expressed NOK in MDA-MB-231 cells, a cell line of triple negative breast cancer (TNBC), a highly metastatic breast cancer type, using an adenovirus (Ad-NOK) (Fig. S1A). Xenograft tumor growth experiments showed that the tumor from Ad-NOK infected cells grew faster than that from Ad-GFP infected cells (Fig. 1D). Consistently, cells infected with Ad-NOK formed larger tumors (Fig. 1E) with heavier tumor weight (Fig 1F) than Ad-GFP infected cells. These results suggest that over-expression of NOK promotes tumor formation and growth for breast cancer cells. Reciprocally, we stably depleted endogenous NOK in MDA-MB-231 cells (Fig. S1B) using two reported siRNAs [39]. Tumor formation assays demonstrated that depletion of NOK significantly reduced the tumor growth rate (Fig. 1G), tumor size (Fig. 1H), and final tumor weight (Fig. 1I) in nude mice. Furthermore, we observed that over-expression of NOK accelerated tumor metastasis in the lung (Fig. 1J-K). Taken together, these results suggest that NOK enhances tumorigenesis and metastasis in TNBC cells.

**Figure 1.**
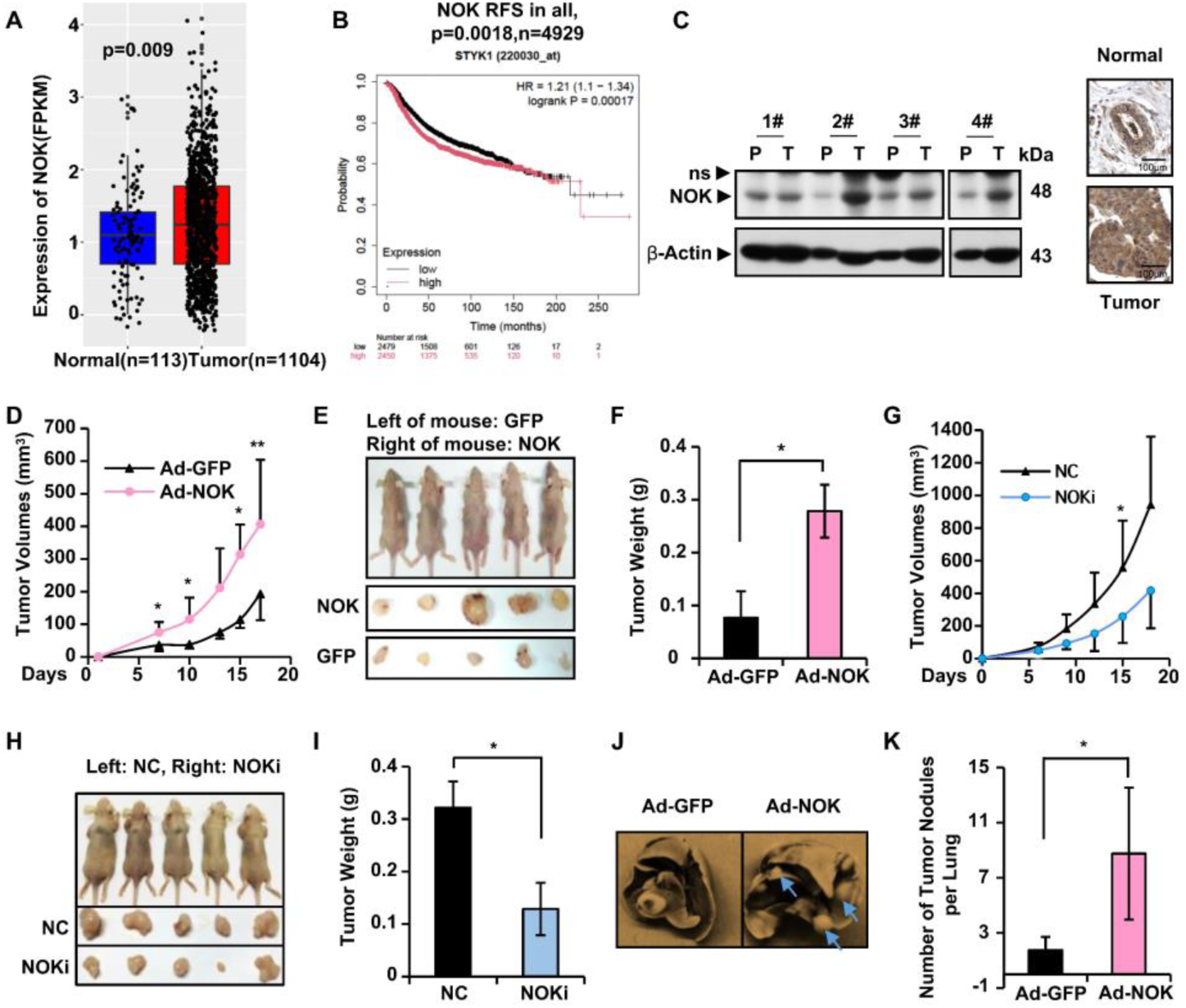
NOK is highly expressed in breast cancer and promotes tumor growth and metastasis of breast cancer cells in nude mice. **A,** Profiles of the expression of NOK/STYK1 in breast tumor and normal tissues. NOK/STYK1 is highly expressed in breast cancers. Boxplots show the mRNA level of NOK/STYK1 in tumor and normal tissues. Data were obtained from TCGA and GTEx database; *p < 0.05. **B**, The expression of NOK predicts the RFS (progression free survival) for the breast cancer patients. A total of 4929 patients were used for the analysis. p = 0.0018. **C**, The protein level of NOK is increased in breast cancers. A western blot was performed for breast cancer samples from 4 patients. P refers to the paired non-tumor tissue and T refers to the tumor tissue from the same patient. β-Actin was used as a loading control. An IHC staining for NOK in breast cancer tumor tissue and paired normal tissue. **D**, Over-expression of NOK enhances the xenograft tumor growth. Nude mice were inoculated subcutaneously with the indicated MDA-MB-231 cells (2 × 10^6^) infected with an adenovirus-driven GFP (Ad-GFP, left arm) and an adenovirus-driven NOK (Ad-NOK, right arm). Tumor volumes at indicated days were calculated by a measurement of width and length in cubic millimeters and presented as mean ± S.D, n = 5, *, *p* < 0.05, **, *p* < 0.01. **E**, A presentation of tumors formed in the nude mice sacrificed at day 18 after the injection of breast cancer cells. Tumors from the left (GFP control) and right (over-expression of NOK) arms in the same mouse are presented. **F**, Over-expression of NOK increases tumor weights. Tumors from the mice were directly weighted. Results are presented as mean ± SD. n=5, *, *p* < 0.05. **G-I**, Depletion of NOK inhibits tumor growth. NOK was depleted in MDA-MB-231 cells by infecting with lentivirus of NOK shRNA. **G**, Tumor volumes from mice inoculated subcutaneously with the indicated cells (5 × 10^6^) were measured in indicated days. Results are presented as mean ± S.D, n = 5, *, *p* < 0.05, **, *p* < 0.01. **H**, A presentation of tumors formed by the mice sacrificed at day 18 after injection of cells. A non-target siRNA was used as a control (NC, on left arm), compared with siRNA-NOK depleted cells (NOKi, right arm). **I**, A quantitative presentation of the tumor weights from NOK depletion cells. Tumors were directly weighed and presented as mean ± SD. n=5, *, *p* < 0.05. **J-K**, NOK promotes metastasis of MDA-MB-231 cells *in vivo*. Equal numbers of MDA-MB-231 cells (5 × 10^6^) infected with Ad-GFP or Ad-NOK were injected intravenously through the tail vein of female nude mice. The whole lung was examined for metastases at 6 weeks after injection. **J**, A demonstration of lungs with metastasis stained with Boulin’s solution. Tumor nodules are in white color. **K**, A quantitative presentation of the tumor nodules in the lung from the nude mice. Values indicate the numbers of tumor nodules as mean ± S.D. *, n=10, *p*< 0.05.

### NOK interacts with EGFR through their juxtamembrane domain (JM) and kinase domain (KD)

As NOK was reported to associate with EGFR, a critical regulator for TNBC, and to co-localize in the early endosome in HeLa cells [43], we determined to study whether NOK promotes breast cancer progression by interacting with EGFR. To this end, we performed immunoprecipitation (IP) experiments in different cells. The results indicated that exogenously expressed EGFR associated with NOK-HA in HEK293T cells (Fig. 2A, * marked bands) and endogenous NOK interacted with EGFR in MDA-MDB-231 cells (Fig. 2B). Consistently, we observed that NOK and EGFR co-localized in the cell membrane and cytoplasm (Fig. S2A), indicating their physical interaction *in vivo*. A FRET analysis by co-expressing YFP-NOK (red, donor) and CFP-EGFR (green, excision acceptor) showed that the interaction between NOK and EGFR occurred in intact living cells (Fig. 2C-2D). Interestingly, we observed that NOK also associated with other members of EGFR family proteins including Her2 (Fig. S2B), Her3 (Fig. S2C) and Her4 (Fig. S2D), implying that NOK might regulate EGFR family receptors. In this study, we focused on the interaction of NOK and EGFR.

**Figure 2.**
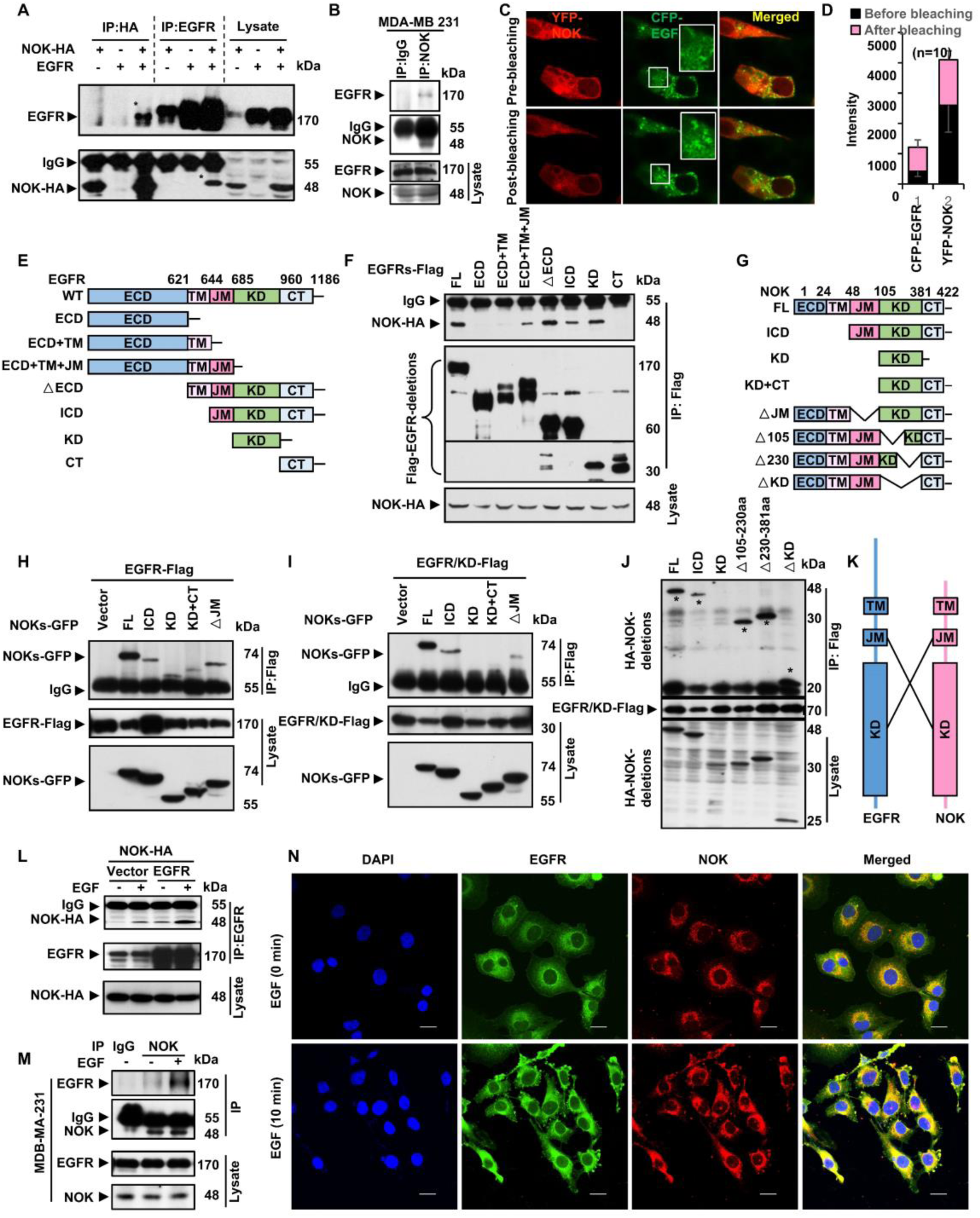
NOK forms a complex with EGFR via an intercross interaction of their juxtamembrane domain and kinase domain. **A**, NOK interacts with EGFR. HA tagged NOK (NOK-HA) was co-expressed with EGFR in HEK293T cells. Cell lysates were reciprocally immunoprecipitated with an antibody against HA or EGFR. The complexes were examined with an antibody against EGFR or HA. The precipitated bands were marked with a star. The protein levels in the cell lysates were showed on right panels. B, Endogenous NOK and EGFR interact in breast cancer cells. An immunoprecipitation (IP) was performed in MDA-MB-231 cells using an antibody against NOK. **C-D**, NOK associates with EGFR in intact cells. **C,** A FRET experiment was performed using HEK293T cells with co-expression of YFP-NOK (red, excision receptor) and CFP-EGFR (green, donor). Images were obtained using a confocal laser scanning microscope. YFP-NOK was photo-bleached and the donor signal (CFP-EGFR) in defined regions of interest (ROI) was measured from at least 10 cells in three different experiments**. D**, The averaged intensities of YFP-NOK and CFP-EGFR before (black column) and after (pink column) bleaching were showed. **E**, A diagraph for different deletions of EGFR. All the mutants were constructed into a Flag-tagged vector. **F**, NOK associates with EGFR through the juxtamembrane domain (JM) and kinase domain (KD) of EGFR. An IP experiment was performed using the cells with co-expression of NOK and EGFR deletions. **G**, A diagraph for deletions of NOK. The mutants were constructed to a GFP-tagged vector. **H**, EGFR associates with NOK via the transmembrane (TM), juxtamembrane (JM) and kinase (KD) domains of NOK. Flag-EGFR was co-expressed with GFP tagged deletions of NOK in HEK293T cells. An IP was performed using an anti-Flag antibody. **I**, The transmembrane domain (TM) and juxtamembrane domain (JM) of NOK are critical for the interaction with the kinase domain of EGFR (KD). Note that the KD of NOK did not interact with the KD of EGFR. The kinase domain of EGFR was co-expressed with deletions of NOK for an IP analysis. **J**, The kinase domain (KD) of NOK had no interaction with the kinase domain of EGFR. Different deletion mutations of NOK and EGFR without its kinase domain (EGFR/KD) were co-expressed in Flag or HA tagged form in 293T cells. A co-IP was performed using an antibody against Flag. The precipitated complex was analyzed by a Western blot using an antibody against HA. **K**, A cartoon to show the cross interaction of the kinase domain of NOK with the TM/JM of EGFR and the kinase domain of EGFR with the TM/JM of NOK. **L-N**, EGF enhances the interaction of NOK with EGFR. **L**, The interaction of HA tagged NOK and EGFR is enhanced by the treatment of EGF (50 ng/ml) at 37°C for 10 min in HEK293T cells. Both NOK and EGFR were over-expressed in HEK293T cells. The immunoprecipitates were analyzed by Western blots. **M**, The interaction of endogenous NOK and EGFR is enhanced by EGF in MDA-MB-231 cells. Immunoprecipitations were performed for endogenous NOK and EGFR proteins. MDA-MB-231 cells were starved for 12 h and maintained in 4°C for 1 h and then treated with EGF at 37°C for 10 min. Cells were harvested and immunoprecipitated with an anti-NOK antibody. The immunoprecipitates were analyzed by Western blots. **N**, Immunostaining assays were performed in MDA-MB-231 cells. Cells were treated as in B and then stained with an anti-EGFR antibody (green), anti-NOK antibody (red) and DAPI (blue).

To examine which domain is critical for the interaction of NOK and EGFR, we constructed deletion mutants of EGFR (Fig. 2E). An IP experiment showed that extracellular domain (ECD) and c-terminus (CT) failed to interact with NOK, whereas the full length, ECD with transmembrane and juxtamembrane (ECD+TM+JM), deletion of extracellular domain (△ECD), intracellular domain (ICD), and kinase domain (KD) mutants maintained the interaction (Fig. 2F). These results suggest that the JM and KD domains of EGFR are critical for the interaction with NOK. Of note, it appeared that TM also contributed to the interaction although ECD+TM showed a very weak interaction (Fig. 2F, lane 4). On the other hand, we mapped domains of NOK (Fig. 2G) in response to the interaction with EGFR. IP experiments showed that the KD domain of NOK exhibited an interaction with EGFR, similar to the ICD, KD+CT and △JM (Fig. 2H). Interestingly, all the interaction between EGFR and KD, ICD or △JM of NOK was weakened (Fig. 2H, lanes 3 and 6), indicating that KD, TM and JM of NOK are critical for the interaction with EGFR. Further IP experiments revealed that the interaction of the kinase domain of EGFR with ICD and △JM of NOK was reduced and no interaction of the kinase domain of EGFR with the KD of NOK was observed (Fig. 2I). Consistently, we observed that deletion of the KD domain (△KD) of NOK had no influence on the interaction with the KD of EGFR (Fig. 2J, * marked), suggesting that the kinase domains of NOK and EGFR are not directly associated when the two receptors form a complex. Taken together, these results suggest that NOK interacts with EGFR mainly via the kinase domain of EGFR and the JM domains of NOK, and EGFR interacts with NOK mainly via the KD of NOK and the JM domains of EGFR. This provides a scenery of the cross interaction structure of the JM domains with the kinase domains of two receptors (Fig. 2K).

### EGF enhances the interaction of NOK with EGFR

To determine whether the interaction of NOK and EGFR is dependent on EGF, a major ligand of EGFR, we performed IP experiments in the cells stimulated with or without EGF. The result showed that the interaction between exogenously expressed NOK and EGFR was enhanced by EGF in HEK293T cells (Fig. 2L). The enhanced interaction between endogenous NOK and EGFR was also observed in MDB-MA-231 cells (Fig. 2M). Consistently, the co-localization of NOK and EGFR was enhanced when EGFR was activated by addition of EGF in MDA-MB-231 (Fig. 2N) breast cancer cells. All the results suggest that the interaction of NOK and EGFR is boosted by EGF.

To further examine whether the interaction of NOK with EGFR is dependent on their phosphorylation, we generated different mutants. A Western blot showed that NOK appeared to preferably associate with EGFR (K721A), a mutant with inactivated kinase, in the absence of EGF, while the interaction with EGFR and EGFR (Y845F), a mutant not activated by c-Src, remained being induced by EGF (Fig. S2E). These results imply that the interaction of NOK and EGFR is dependent on the relaxed conformation of EGFR. Reciprocally, we used two NOK mutants (Y356F and Y327F) that lost the ability of tumor formation [14], and a constitutively active mutant Y417F [38]. The results showed that these mutants remained interacting with EGFR although the interaction of EGFR with NOK (Y327F) and NOK (Y356F) was slightly decreased (Fig. S2F). All these results suggest that the interaction of NOK and EGFR is dependent on EGF but not on the kinase activity of EGFR and NOK.

### NOK and EGFR promote tumorigenesis synergistically

We questioned whether NOK and EGFR coordinately regulate breast cancer cell growth. For this purpose, we examined cell proliferation following over-expression or depletion of NOK in the presence of EGF. Intriguingly, we observed that over-expression of NOK dramatically promoted MDA-MB-231 cell growth in the presence of EGF (Fig. 3A, 72 h, blue vs red column). Reciprocally, depletion of NOK by a mixture of siRNAs abolished the effect of EGF on cell proliferation (Fig. 3B, compare red and blue columns at 96 h). Soft agar assays showed that NOK promoted MDA-MB-231 cells to form more colonies in the presence of EGF (Fig. S3A, S3B). Consistently, we observed that over-expression of NOK significantly increased migration of MDA-MB-231 cells in the presence of EGF (Fig. 3C-4D, S3C-S3D), while depletion of NOK dampened EGF-induced cell migration (Fig. 3E-F). These results suggest that NOK coordinately activates the EGF signal for breast tumor cell proliferation and migration. To illustrate the role of NOK in coordination with EGF, we used MEFs from NOK knock-out mice (NOK KO) (Fig. S3E). Cell proliferation assays showed that EGF promoted cell growth of wild type MEFs but not NOK KO MEFs (Fig. 3G). These results confirmed that NOK is essential for EGF on cell proliferation.

**Figure 3.**
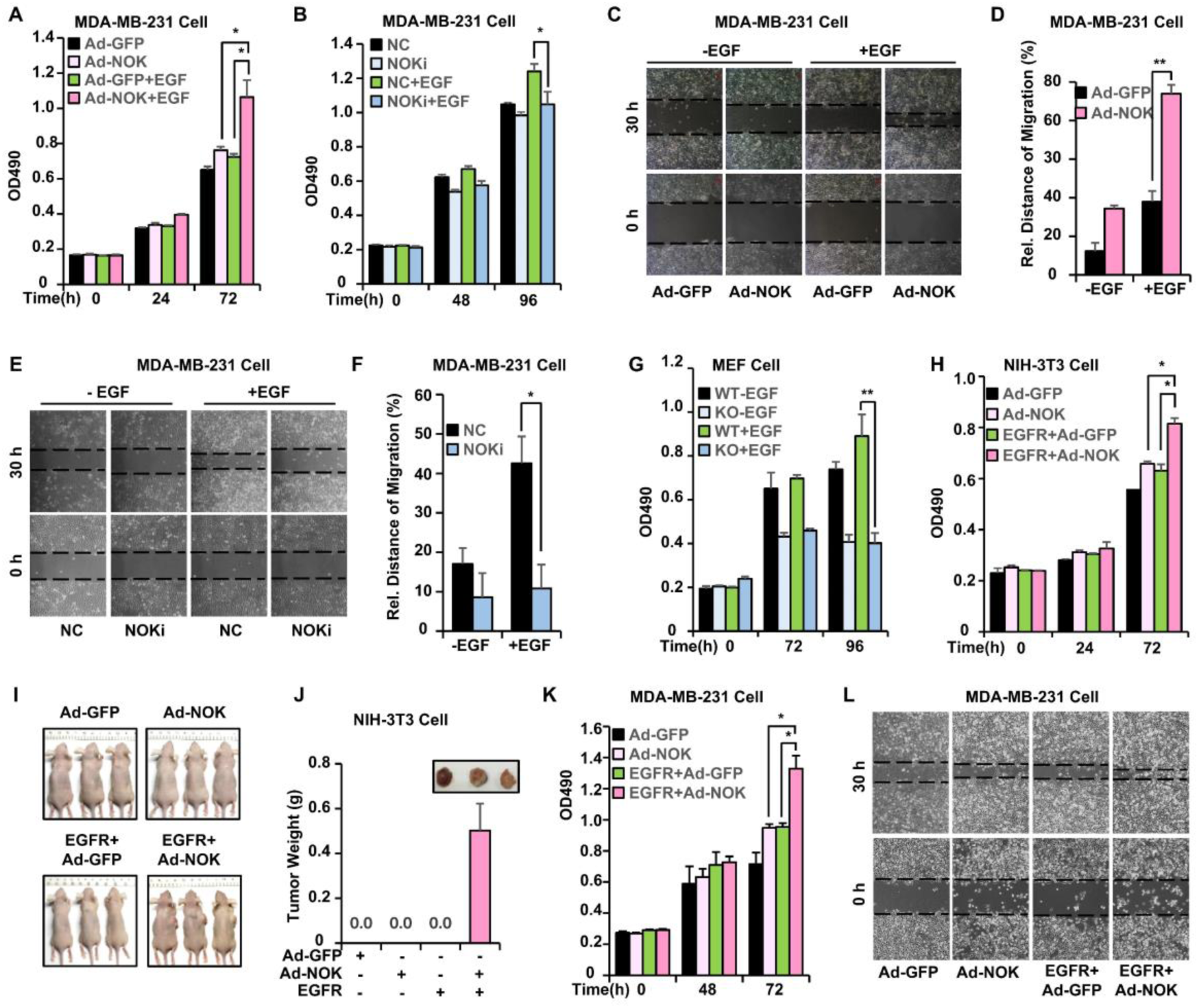
NOK synergistically reinforces EGF-induced breast cancer cell proliferation and migration. **A**, NOK enhances the EGF-induced cell proliferation. MDA-MB-231 cells were infected with Ad-NOK or Ad-GFP and seeded in triplicate on 96-well plates (1 × 10^3^/well), treated with or without EGF (50 ng/ml) for the indicated times. Cell numbers were measured at OD 490 nm/630 nm. **B**, EGF-induced cell proliferation is impaired by NOK depletion. MDA-MB-231 cells were infected with a lentivirus expressing control shRNA (NC) or NOK specific shRNA (NOKi) were seeded in triplicate on 96-well plates (1.2 × 10^3^/well). Assays were performed as in A. **C-D**, NOK enhances the EGF-induced cell migration. **C**, MDA-MB-231 cells were infected with Ad-NOK or Ad-GFP. Cells were seeded in triplicate on 6-well plates (5 × 10^4^/well) for the wound healing assay. Confluent monolayer cells were scraped with sterile 200 ul pipette tip and allowed to migrate for 30 h in the presence or absence of EGF (50 ng/ml). The wounded areas were photographed at 0 and 30 h. **D**, Relative migration rates were measured by Image J and cell migration distances are presented as means ± S.D. *, *p*< 0.05. **E-F**, Depletion of NOK impairs cell migration mediated by EGF. MDA-MB-231 cells were infected with a lentivirus expressing control sh RNA (NC) or NOK specific shRNA (NOKi). Confluent monolayer cells were scraped with sterile 10 ul pipette tip and assays were performed as in C-D. **G**, MEF cells from wild type (WT) and NOK knocked-out mice (KO) were seeded into triplicate on 96-well plates (1.0 × 10^3^/well) and treated with or without EGF (50 ng/ml) for indicated times. Cell numbers were measured at OD 490 nm/630 nm. **H**, Over-expression of NOK and EGFR synergistically promotes cell proliferation. EGFR was stably over-expressed by a lentivirus and NOK was over-expressed by an adenovirus (Ad-NOK). Ad-GFP was used as a control. NOK and EGFR have a synergistic effect on cell proliferation induced by EGF. The NIH-3T3 cells expressing different combinations of NOK and EGFR were seeded in triplicate on 96-well plate (1 × 10^3^/well) and treated with or without EGF (50 ng/ml) for indicated time periods. Cell numbers were measured at OD 490 nm/630 nm. **I-J,** NOK and EGFR have a synergistic effect on the tumorigenesis. NIH-3T3 cells with over-expression of NOK and/or EGFR were implanted in nude mice for the tumorigenesis. Mice are showed after 8 weeks after implantation of the cells (I). Tumors from the mice were directly weighted (J). Results are presented as mean ± SD. n=3. Note that no tumors were observed in the mice implanted with cells with Ad-GFP, Ad-NOK or EGFR. **K**, NOK and EGFR synergistically promote the proliferation of MDA-MB-231 cells. EGFR was stably over-expressed and NOK was over-expressed by an adenovirus (Ad-NOK). Ad-GFP was used as a control. The MDA-MB-231 cells expressing different combinations of NOK and EGFR were seeded in triplicate on 96-well plate (1 × 10^3^/well) and treated with or without EGF (50 ng/ml) for indicated time periods. Cell numbers were measured at OD 490 nm/630 nm. *, *p*< 0.05 **L**, NOK and EGFR have a synergistic effect on cell migration induced by EGF in the MDA-MB-231 cells. Assays were performed as in C and E and cells were used as in K.

**Figure 4.**
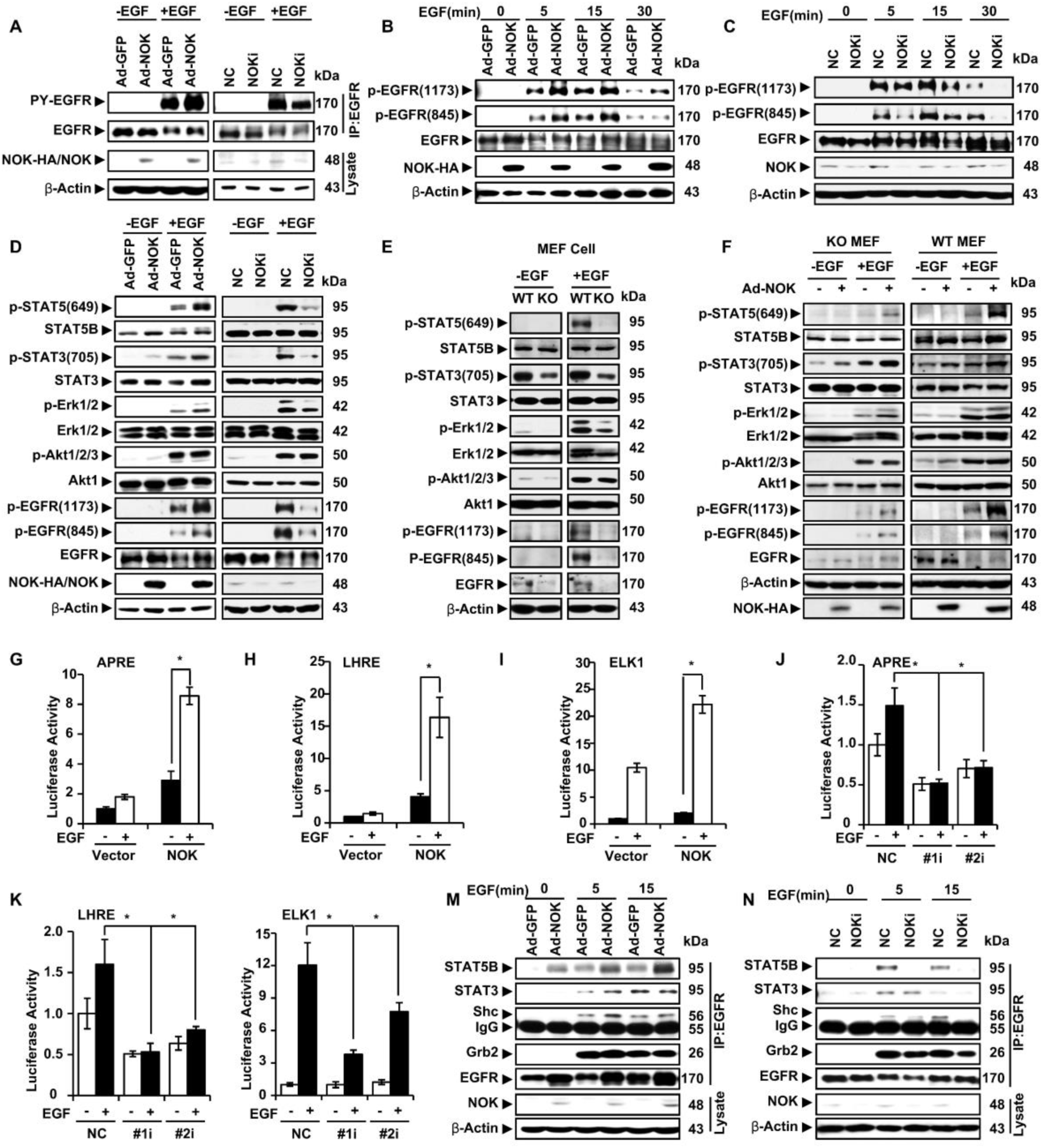
NOK enhances EGFR activation and downstream signaling. **A**, NOK enhances the tyrosine phosphorylation of EGFR. NOK was over-expressed by an adenovirus (Ad-NOK, left panel) or depleted by transfection of an siRNA (NOKi, right panel) in MDA-MB-231 cells. An adenovirus expressing GFP (Ad-GFP) and a nonspecific siRNA (NC) were used as negative controls in the experiment. Cells were treated with or without EGF (100 ng/ml) for 5 min after starvation for overnight. Phosphorylated EGFR (pY-EGFR) was examined with a pY(4G10) antibody from an EGFR-complex precipitated by an antibody against EGFR (sc-120). Total EGFR levels were examined with an EGFR antibody (sc-03). **B-C**, NOK promotes the phosphorylation of EGFR in response to EGF within 30 min. NOK was over-expressed (B) or depleted (C) in MDA-MB-231 cells treated with or without EGF (100 ng/ml) for the indicated times. Phosphorylated EGFR was examined with phosphorylation antibodies against two different tyrosine residues (Y1173 and Y845). **D,** NOK enhances the activation of EGFR to phosphorylate the downstream signaling effectors. NOK was over-expressed (left panel) or depleted (right panel) in MDA-MB-231 cells treated with or without EGF (100 ng/ml) for 5 min. Phosphorylated and total proteins of STAT3, STAT5B, Erk1/2 and Akt1/2/3 were examined with the antibodies as indicated. NC refers a non-specific siRNA comparing to an siRNA against NOK (NOKi). **E**, The activation of EGFR signaling was impaired when NOK was knocked out. MEF cells from wild type (WT) or NOK knock-out (KO) mice were cultured in the presence or absence of EGF (100 ng/ml) for 5 min (left panel). **F**, Downstream effectors of EGFR signaling were recovered by exogenous expression of HA-NOK in NOK KO mice (right panel). The phosphorylation of STAT3, STAT5B, Erk1/2 and Akt1/2/3, as well as EGFR, were examined. **G-I**, NOK up-regulates the transcriptional activity of STAT3, STAT5, and ELK1 in response to EGF signaling. Over-expression of NOK (G-I) promotes the activation of STAT3 (APRE-luciferase), STAT5 (LHRE-luciferase), and ELK1 in the presence of EGF stimulation. The luciferase reporters were co-transfected respectively with pRL-TK plasmids into MDA-MB-231 cells with over-expression (G-I) of NOK. The transcriptional activity was expressed as fold-changes, normalized by an internal control (Renilla). Results were from three independent repeats and are presented as mean± S.D. *, *p*<0.05. **J-L**, Depletion of NOK reduces the transcriptional activity of STAT3, STAT5, and ELK1 in response to EGF signaling. NOK was depleted together with the expression of STAT3-, STAT5-, and ELK1-driven luciferase reports in the presence of EGF stimulation. The luciferase experiments were performed as in G-I. **M-N**, NOK promotes the recruitment of effector proteins to EGFR in response to EGF stimulation. NOK was over-expressed (M) or depleted (N) in MDA-MB-231 cells treated with EGF (100 ng/ml) for indicated times. The complex of EGFR was deciphered by Western blots from a precipitant of an antibody against EGFR (IP: EGFR). The EGFR complex contains STAT3, STAT5B, Shc and Grb2 dissected with indicated antibodies. β-Actin was used as a loading control in all the experiments.

To examine whether NOK and EGFR have a synergistic effect on cell growth, we over-expressed EGFR and NOK in NIH-3T3 cells, a normal fibroblast cell line without endogenous EGFR. Cell proliferation assays indicated that co-expression of EGFR and NOK had a significant synergistic effect on cell growth in comparison with either control or expression of EGFR or NOK in NIH-3T3 (Fig. 3H). Intriguingly, we observed that co-overexpression of NOK and EGFR transformed the NIH-3T3 cells to form tumors (Fig. 3I-4J). This result suggests that NOK is required for EGFR during its induction of tumorigenesis. Simultaneously, we observed that over-expression of NOK dramatically increased the effect of EGFR on cell proliferation and migration in MDA-MB-231 cells (Fig. 3J). Taken together, these results suggest that NOK and EGFR function synergistically on tumorigenesis.

### NOK enhances EGF signaling mainly on its downstream effectors STAT3, STAT5, and Erk1/2

To reveal how NOK enhances EGF signaling on tumorigenesis, we investigated the phosphorylation of EGFR under over-expression or depletion of NOK in MDA-MB-231 cells. Western blot analyses showed that over-expression of NOK and EGFR in MDA-MB-231 (Fig. S4A) and NIH-3T3 cells (Fig. S4B) promoted the phosphorylation of EGFR (p-Y1173). Consistently, transient expression of NOK and EGFR in HEK293T cells resulted in enhanced pan-phosphorylation of EGFR (4G10) (Fig. S4C). Intriguingly, we observed that over-expression of NOK enhanced, but depletion of NOK (Fig. S4D) reduced, the phosphorylation level of endogenous EGFR in the presence of EGF (Fig. 4A). Furthermore, we observed that over-expression of NOK promoted (Fig. 4B), but depletion of NOK inhibited (Fig. 4C), the phosphorylation of EGFR by the addition of EGF within 30 min. Taken together, these results suggest that NOK regulates phosphorylation of EGFR.

Activation of EGFR leads to the phosphorylation of downstream effectors including STAT3, STAT5, Erk and Akt [1, 5]. To reveal the role of NOK on these EGFR downstream effectors, we examined their phosphorylation status. Western blot analyses indicated that over-expression of NOK enhanced (Fig. 4D, left panels), but depletion of NOK impaired (Fig. 4D, right panels), the phosphorylation of EGFR and downstream STAT3, STAT5, and Erk1/2 significantly, but exhibited little effect on the phosphorylation of Akt1/2/3, in the presence of EGF (Fig. 4D, bottom lanes). Furthermore, we examined the EGFR signaling in wild type and NOK KO MEFs. The results demonstrated that EGF failed to activate the phosphorylation of STAT3, STAT5, and Erk1/2 in NOK KO MEFs (Fig. 4E, left panel), but had no effect on the phosphorylation of Akt1/2/3 (Fig. 4E). These results consistently suggest that NOK is responsible mainly for the phosphorylation of STAT3,

STAT5, and Erk1/2. To validate the role of NOK on EGF-induced phosphorylation of EGFR downstream effectors, we performed a rescue experiment using Ad-NOK (Fig. 4F). Indeed, we observed that the role of EGF on the phosphorylation of EGFR, STAT3, STAT5 and Erk1/2 was recovered by Ad-NOK in NOK KO MEFs (Fig. 4F, right 2 lanes). Taken together, these results indicate that NOK is essential for the activation of EGFR and the downstream signaling, preferably in promoting the phosphorylation of STAT3, STAT5 and Erk1/2.

To further interrogate the function of NOK on STAT3, STAT5, and Erk1/2, we used luciferase reporters driven by APRE (in response to STAT3), LHRE (in response to STAT5), ELK1 (in response to Erk1/2). The results showed that over-expression of NOK significantly promoted the luciferase activities in the presence of EGF in response to STAT3 (Fig. 4G), STAT5 (Fig. 4H), and Erk1/2 (Fig. 4I) in MDB-MA-231cells. Reciprocally, we observed that the luciferase activities were dramatically damped when NOK was depleted by siRNAs (Fig. 4J-5L). These results suggest that NOK coordinates with EGFR to activate STAT3, STAT5, and Erk1/2, downstream of EGF signaling.

**Figure 5.**
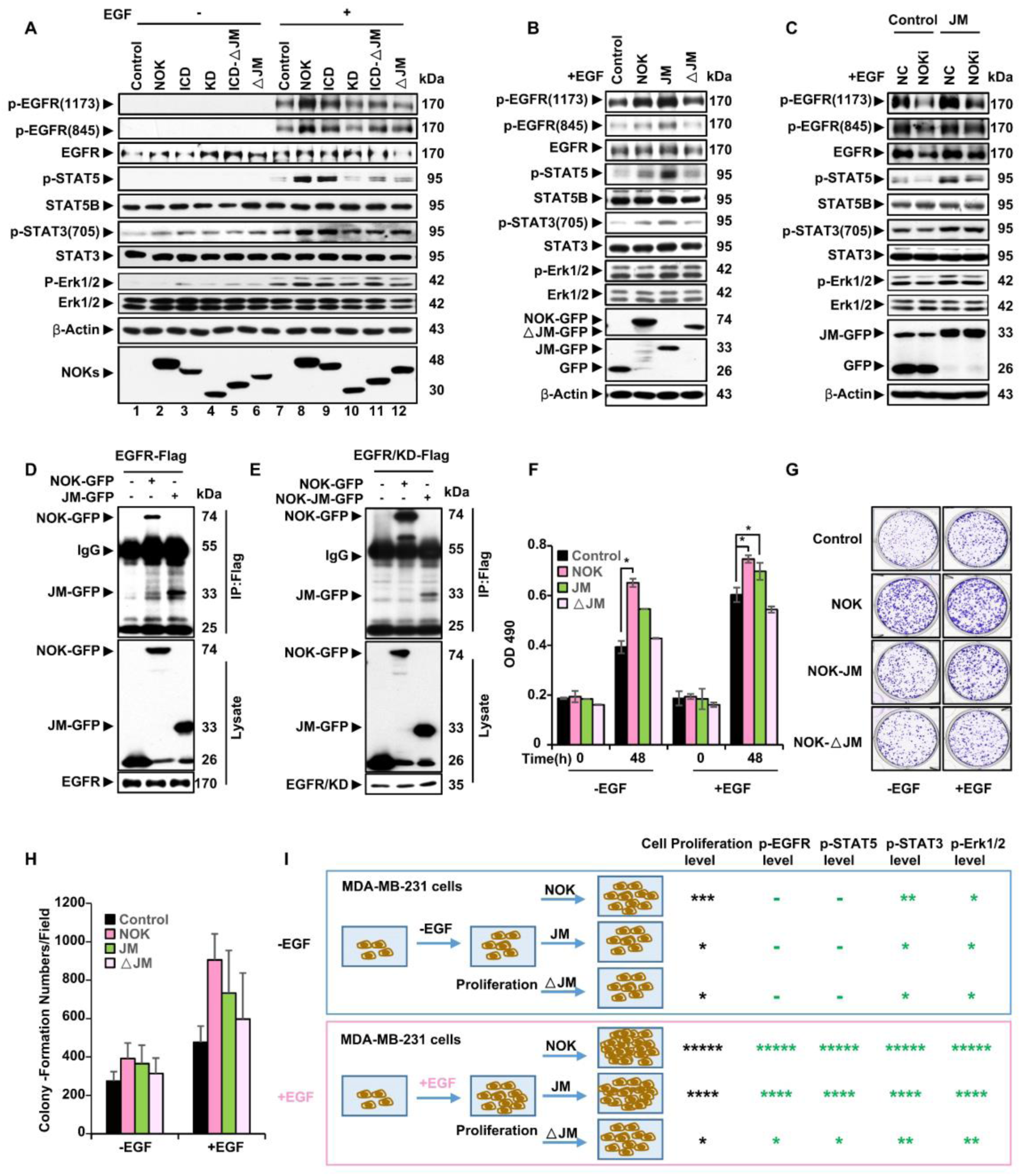
JM domain of NOK mediates EGFR signaling activation. **A,** Depletion of juxtamembrane domain (JM) of NOK (ΔJM) impairs EGFR signaling. Deletions of NOK were expressed in MDA-MB-231 cells treated with or without EGF (100 ng/ml) for 5 min. The phosphorylation of EGFR, STAT3, STAT5, and Erk1/2 was examined. **B**, JM of NOK is critical to enhance EGFR signaling. Full length, JM of NOK and ΔJM were expressed in MDA-MB-231 cells. EGFR signaling was examined using the indicated antibodies. Note that JM enhanced, but ΔJM failed to promote, the phosphorylation of EGFR, STAT3, STAT5, and Erk1/2. **C**, JM of NOK partially rescues the activation of EGFR signaling without endogenous NOK. The endogenous NOK protein was depleted by a mixture of siRNAs against NOK (NOKi). Activation of EGFR signaling was examined. A non-specific siRNA (NC) was used as a negative control. **D-E**, JM of NOK associates with EGFR (D) and the kinase domain of EGFR (E). Flag tagged EGFR and the kinase domain (KD) were co-expressed with GFP tagged full length and JM of NOK for IP experiments using an antibody against Flag. **F-H**, JM of NOK enhances the proliferation (F) and colony formation (G) in breast cancer cells. MDA-MB-231 cells (1 × 10^3^) were stably transfected with plasmids of control vector, NOK, JM and ΔJM. Cell densities (F) were measured at OD 490 nm/630 nm after EGF-treatment (for 48 h) of cells cultured in 96-well plates. Results of colony formation were showed in G. **H,** The colonies in the whole field (Fig. 5G) were counted. Values indicate the numbers of colonies as mean ± S.D. *, p<0.05, n=3. **I,** A graphic model of NOK, NOK-JM, NOK-ΔJM in mediating the EGFR signaling and cell proliferation. NOK and NOK-JM promote the EGF signaling, cell proliferation and colony formation under the EGF stimulation.

As activation of EGF signaling occurs upon the recruitment of adaptor proteins to EGFR, we examined whether NOK promotes the complex formation of EGFR with STAT3, STAT5, Shc and Grb2. IP experiments using an antibody against EGFR revealed that both STAT3 and STAT5B existed in the EGFR complex, together with Shc and Grb2 (Fig. 4M). Of note, over-expression of NOK promoted the complex formation at different time points after EGF treatment (Fig. 4M, see 5, 15 min). Simultaneously, we observed that depletion of NOK decreased the recruitment of STAT3, STAT5B, Shc and Grb2 (Fig. 4N). These results suggest that the recruitment of all these four proteins is mediated by EGF and is enhanced by NOK. Therefore, we conclude that NOK promotes the EGF-induced activation of EGFR downstream effectors by recruiting STAT3, STAT5B, Shc and Grb2 to EGFR.

### NOK mediates activation of EGFR through its JM domain

Since both of the JM and TM domains of NOK are critical for its interaction with EGFR, we questioned whether these domains are critical for the activation of EGFR. To this end, we examined the phosphorylation of EGFR downstream effectors in the presence of different NOK mutants in MDB-MA-231 cells. A Western blot analysis demonstrated that over-expression of NOK significantly increased the phosphorylation of EGFR, as well as activation of STAT3, STAT5, and Erk1/2 (Fig. 5A, comparing lane 8 to 7). Interestingly, deletion of JM failed to enhance the phosphorylation of EGFR, STAT3, STAT5, and Erk1/2 (Fig. 5A, lane 12), but deletion of TM (ICD) had no significant effect on the phosphorylation of these proteins (Fig. 5A, lane 9). This result suggest that although TM, JM and KD domains of NOK remained the interaction with EGFR, the JM domain appeared to be critical for the activation of EGFR. To examine the role of JM domain on the EGFR downstream effectors, we over-expressed the JM domain of NOK in cells treated with EGF. Western blot results showed that expression of JM alone strongly enhanced, but deletion of JM failed to promote, the phosphorylation of EGFR, STAT3, STAT5, and Erk1/2 (Fig. 5B). Of note, among these effectors, STAT5 phosphorylation was the most significantly affected by NOK (Fig. 5A-6B, also see Fig. 4). These results suggest that the JM domain of NOK is critical for the activation of EGFR and its downstream effectors, in particular STAT5.

**Figure 6.**
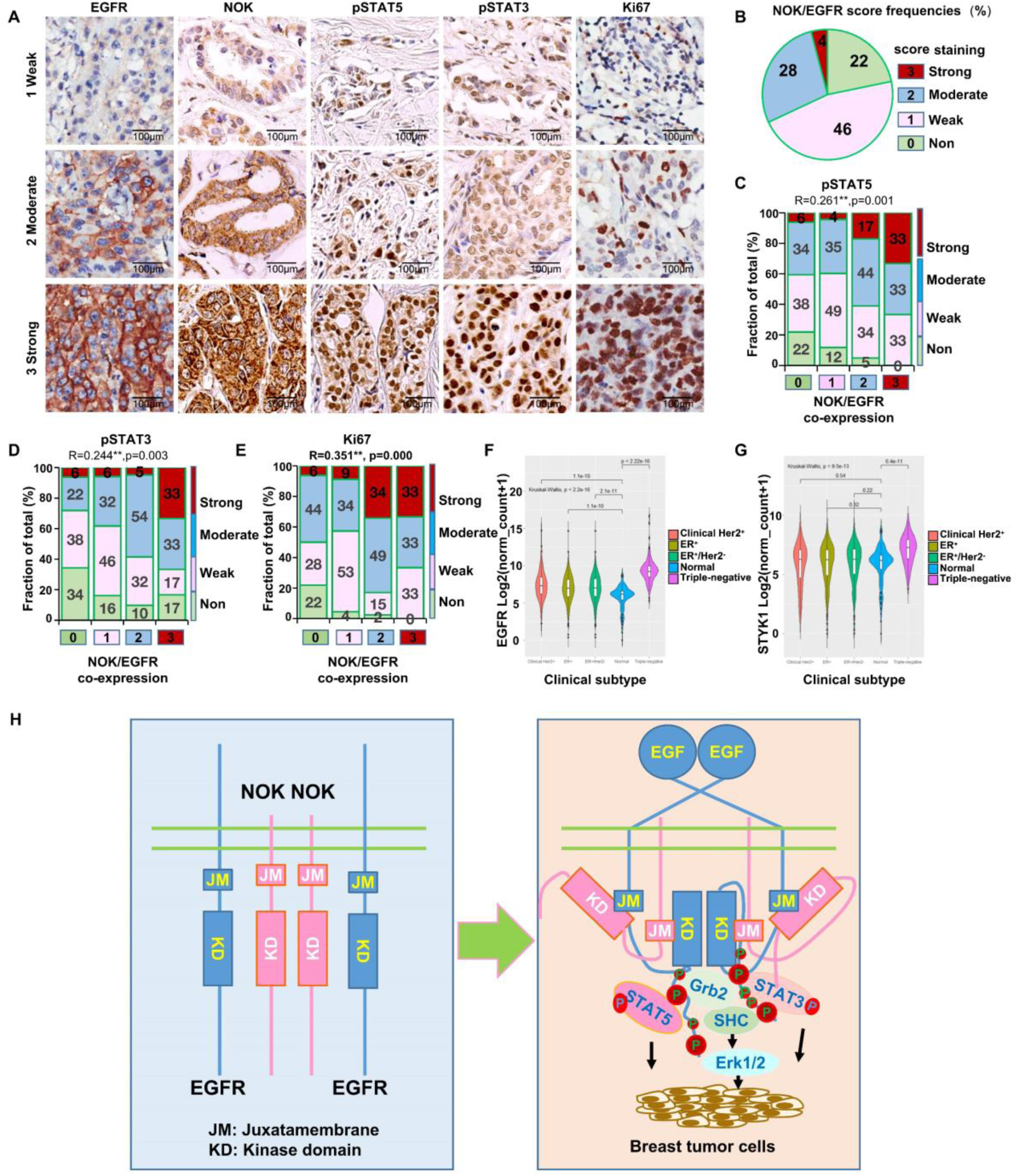
Co-expression of NOK and EGFR is correlated with the expression of p-STAT5, p-STAT3, and Ki67 in breast cancer tissues. **A**, The expression levels of EGFR, NOK, pSTAT5, pSTAT3, and Ki67 were presented as weak, moderate and strong in human breast adenocarcinoma. Consecutive sections of resected human breast adenocarcinomas were stained for the indicated proteins. **B**, The frequency of NOK and EGFR co-expression in breast cancers. The co-expression of NOK and EGFR was defined in 4 grades as strong, moderate, weak and none, based on their individual expression. The score was calculated as minimized scores of EGFR and NOK. **C-D**, Co-expression of NOK and EGFR correlates the phosphorylation of STAT5 and STAT3 in different stages of breast cancers. Case numbers were labeled for each grades and correlation efficiency was marked with p value. ** indicated significant level. **E**, Co-expression of NOK and EGFR also correlates to the level of Ki67, an indicator for the tumor cell proliferation ability. A total number of 147 patients were used. **F-G**, Gene expression profiles of EGFR and NOK in different types of breast cancer. TCGA Breast Cancer (BRCA) (24 datasets) were downloaded from publicly available website (https://xena.ucsc.edu/) in this study. This dataset shows the gene-level transcription estimates, as in log2(x+1) transformed RSEM normalized count. All data were processed and analyzed by Excel 2010 and R (version 4.3.3). **H,** A graphic model of NOK and EGFR in mediating the signaling transduction during tumorigenesis. NOK and EGFR form heterodimers under the EGF stimulation. NOK promotes the EGFR phosphorylation through the NOK JM domain and the EGFR kinase domain interaction. The association of NOK and EGFR activates the EGFR downstream effectors STAT3, STAT5, and Erk1/2 to facilitate the tumor transformation of normal cells and to accelerate the tumor cell proliferation and migration. NOK and EGFR synergistically enhance the tumor growth and metastasis of breast cancer cells.

To further confirm the role of JM in the activation of EGFR, we depleted endogenous NOK and re-expressed its JM domain. A Western blot showed that exogenous JM partially rescued the function of NOK on EGFR signaling (Fig. 5C). This result suggests that the JM domain of NOK plays a critical role in the activation of EGFR, consistent with our aforementioned observations on the critical role of JM for the interaction with EGFR (Fig. 2 and S2). We also confirmed that the JM domain of NOK alone interacted with EGFR (Fig. 5D) and the KD domain of EGFR (Fig. 5E). Therefore, we conclude that NOK functions on EGFR via its JM domain interacts with the KD of EGFR..

To validate the function of JM domain, cell proliferation and colony formation assays were performed using GFP, GFP-NOK, GFP-JM and GFP-ΔJM, which were stably over-expressed in MDA-MB-231 cells (Fig. S5A). The result showed that in the absence of EGF, the growth of cells stably expressing NOK was the fastest among the four cell lines, followed by JM over-expressing cells, but cells with deletion of JM of NOK grew at the same rate as the control cells (Fig. 5F. left). In the presence of EGF, NOK and JM expressing cells grew faster than control cells and JM deletion cells (Fig. 5F, right). Consistently, cells expressing NOK and JM formed more colonies than control cells and ΔJM expressing cells in the presence of EGF (Fig. 5G-H). All these results indicate that the JM domain is critical for the activation of EGF signaling in regulating cell proliferation and colony formation (Fig. 5I).

### Co-expression of NOK and EGFR is correlated with p-STAT5, p-STAT3 and Ki67 levels in breast cancer tissues

To investigate whether the expression of NOK and EGFR is related to human breast cancers, we focused on STAT3/5 and performed immunostaining using human breast cancer samples. We classified the staining density as 4 grades ranging from 0 to 3 for NOK, EGFR, pSTAT5, pSTAT3 and Ki67, a cell proliferation marker (Fig. 6A). Interestingly, we observed that NOK and EGFR were co-expressed in 78% of the tumors, of which 28% with high and 4% with strong co-expression (Fig. 6B). Since our aforementioned results demonstrated that NOK and EGFR synergistically promoted phosphorylation of STAT3 and STAT5, we questioned whether co-expression of NOK and EGFR correlates with the levels of p-STAT3 and p-STAT5 in human cancers. We found that strong co-expression of NOK and EGFR correlated to high levels of p-STAT5 (Fig. 6C), p-STAT3 (Fig. 6D), and Ki67 (Fig. 6E). Using Pearson correlation analysis, we found that the expression of NOK correlated with levels of p-STAT5 (R=0.349, p<0.01), p-STAT3 (R=0.316, p<0.01), and Ki67 (R=0.439, p<0.01), however, expression of EGFR alone only correlated with the level of Ki67 (R=0.285, p<0.01), but not with the levels of p-STAT5 or p-STAT3 (Table S1). Intriguingly, a Pearson correlation analysis indicated that co-expression of NOK and EGFR correlated with levels of pSTAT5, pSTAT3 and Ki67 (R=0.261, 0.244 and 0.351, respectively, all p<0.01) (Table S2).

Finally, we analyzed the expression of NOK and EGFR in breast cancer patients from a public website (https://xena.ucsc.edu/). The data from the TCGA Breast Cancer (BRCA) (24 datasets) were downloaded and analyzed in this study. The results showed that both NOK and EGFR were highly expressed in different types of breast cancers, in particular in the TNBC patients (Fig. 6F-7G). All these results suggest that NOK and EGFR coordinately regulate tumorigenesis in breast cancers.

## DISCUSSION

Over-activation of EGFR signaling is closely related with tumorigenesis [1]. In the present study, we found that NOK, a tyrosine receptor-like transmembrane protein, is a co-activator of EGFR to promote tumorigenesis. We revealed that NOK is essential for the activation of EGFR via its JM domain interacting with the kinase domain of EGFR (Fig. 8). Intriguingly, NOK-associated EGFR appears to preferably phosphorylate downstream effectors STAT3 and STAT5 in breast cancer cells. Our study confirmed the interaction of NOK with EGFR as reported by others [32–34] and extended the role of NOK on the activation of EGFR. Importantly, we found that NOK and EGFR synergistically promote cell transformation and tumor formation, indicating that NOK is critical for EGF-induced breast cancers. At the clinical level, we observed that NOK and EGFR are co-expressed and strongly correlated with Ki67, a proliferating marker for tumor growth, and also the levels of p-STAT3 and p-STAT5, two critical factors in promoting tumor malignance, in human breast cancer tissues. Our study reveals an axis of NOK-EGFR-activated STAT3 and STAT5 signaling during the tumorigenesis of breast cancers. We speculate that targeting the JM domain of NOK will benefit the therapeutic effect among breast cancer patients with high EGFR expression. As EGFR signaling is extensively studied in different cancers, we postulate that NOK-regulated EGFR activation may also participate in other cancers.

Previous studies indicated that EGFR signaling requires an asymmetric interaction and trans-activation between the kinase domains of two receptors [4, 44–46]. The kinase domain of one receptor molecule functions as the activator to awaken the kinase domain of the other receptor, which is regarded as the receiver [44, 45, 47]. Formation of the asymmetric dimer appears to underlie the activation of all EGFR family members [44]. The intracellular JM segment of the receptor, known to potentiate the kinase activity, is able to dimerize the kinase domains [45]. In this study, we observed that the JM domain of NOK interacted with the kinase domain of EGFR, resembling the role of the EGFR-JM domain, where the JM domain of receiver EGFR latches the kinase domain of receiver EGFR to the activator EGFR [45]. Therefore, we speculate that NOK may function in a similar way as the receiver EGFR, where the JM domain of NOK latches the activated kinase domain to the activator EGFR. This asymmetric dimer pattern defines a specific event to phosphorylate STAT3, STAT5 and Erk1/2, downstream of EGFR. Our data support the model that NOK plays a role in unravelling the kinase domain of EGFR in the NOK-EGFR heterodimer and simultaneously recruiting STAT3 and STAT5, also possibly other activator of Erk1/2. This is similar to the role of Her2 in the hetero-dimer with EGFR, although the dimer formation of EGFR with Her2 is mediated by the kinase domains of two receptors, whereas the dimer of EGFR and NOK is maintained by the JM domain of NOK and the kinase domain of EGFR [48]. However, we could not exclude other possibilities. For instance, it is also possible that NOK may form tetramer with EGFR to stabilize asymmetric dimer of EGFR. This hetero-tetramer seems to form through their TMs [49], as we observed that the intracellular domain of NOK (ICD) had a very weak interaction with EGFR and the TM domain was critical for their interaction (Fig. 2I). This may provide an opportunity for EGFR to recruit different effectors. In such a way, NOK exhibits a diverse role for the regulation of EGFR activation, however, detailed molecular mechanism underlying such a process requires further studies.

Accumulating evidence demonstrated that the EGFR activity could be regulated at different levels, including dimerization (e.g. Mig6, Cytohesins) [50, 51], protein stability (e.g. PTK6, TNK2) [52, 53], protein modification (e.g. HDAC6, HDAC-3/CAGE Axis) [54, 55], internalization (e.g. Mig6), and the kinase activity (e.g. Mig6) [56, 57]. Our study provides strong evidence that NOK dimerizes with EGFR to activate EGFR signaling by promoting EGFR kinase activity. However, it remains unclear whether the NOK-EGFR dimer functions to stabilize EGFR. Basically, dimerized EGFR undergoes internalization and then is degraded by the lysosome [54]. The internalization process of EGFR is regulated by Rab family proteins, which mediates the sorting of the internalized EGFRs [1]. Whether NOK regulates the internalization of EGFR is of interest for future study.

EGFR has been reported to activate different downstream pathways. In this study, we observed that NOK enhanced the activation of EGFR to activate STAT3 and STAT5 as well as Shc and Grb2, which correspond to Erk1/2 activation. Our results demonstrated that the interaction of NOK with EGFR specifies EGFR signaling to STAT3, STAT5, and Erk1/2 but not Akt. This seems quite different from its role on GSK-3beta, where NOK associates with Akt and GSK-3beta [58]. We speculate that NOK may associate with different effectors in different cells. In this study, we presented that the main function of NOK is to facilitate EGFR to activate STAT3, STAT5, and Erk1/2, leading to cell proliferation and migration in breast cancer cells. Our results also echo the observations from the expression patterns of EGFR and phosphorylated STAT3 and STAT5 in pathological studies. Therefore, we conclude that NOK defines a specific activation pathway for EGFR to activate STAT3, STAT5, and Erk1/2 during the tumorigenesis of breast cancers.

We proposed that NOK is a co-receptor for EGFR and focused on the effect of NOK on the phosphorylation of EGFR. Reciprocally, it is also possible that EGFR is a co-receptor for NOK. A recent study reported that EGFR phosphorylated NOK at the Y356 residue via the interaction between the kinase domains [32]. Indeed, our previous study demonstrated that NOK interacted with c-Src via the kinase domains of the two proteins [31]. However, in this study, we observed that EGFR interacted with NOK via a cross association of TM/JM and kinase domains of these two proteins. Although the interaction of EGFR and NOK was slightly weakened when the residue Y356 in the NOK kinase domain was mutated (see Fig. S3B), deletion of the whole kinase domain of NOK remained a strong interaction with EGFR (Fig. S2E). Our results are in consistency with observations that the JM domain in EGFR is critical for the interaction with the kinase domain of receiver EGFR [45, 59]. On the other hand, NOK contains only 22 amino acids in its extracellular domain and no ligand is reported. At this context, we consider that NOK functions as a co-receptor for EGFR mainly via their interaction of the intracellular structures (Fig. 6H).

## CONCLUSION

In the present study, we found that NOK, a tyrosine receptor-like transmembrane protein, is a co-activator of EGFR to promote tumorigenesis. We revealed that NOK is essential for the activation of EGFR via its JM domain interacting with the kinase domain of EGFR. We conclude that NOK defines a specific activation pathway for EGFR to activate STAT3, STAT5, and Erk1/2 during the tumorigenesis of breast cancers. Taking together, our study shows that NOK is a potential EGFR regulator and targeting their interaction providing a new strategy for the therapy of breast cancers.

## ACKNOWLEDGMENTS

We thank Dr. Michael Famulok from 1LIMES Institute, Rheinische Friedrich-Wilhelms-Universitat Bonn, Germany, and Dr. Jose Antonio Rodriguez from Vrije Univ. Amsterdam Med Ctr, Netherlands, to kindly provide EGFR related plasmids.

## AUTHOR CONTRIBUTIONS

Yinyin Wang performed most of the experiments, analyzed the data and wrote the paper. Bingdong Zhang and Chunhua He analyzed part of the data. Bo Tian, Sihan Liu, Jianghua Li, Jiayu Wang, Shigao Yang and Bingtao Zhu were involved in performing the experiments of tumor assay and bioinformatics analyses. Xiaoguang Wang, Zhijie Chang and Chenxi Cao supervised the project, critically reviewed the results and revised the paper.

## ADDITIONAL INFORMATION

### Ethics approval and consent to participate

The experimental protocol was developed in accordance with the ethical guidelines of the Helsinki Declaration. Ethical approval was granted by the Human Ethics Committee. Written informed consent was obtained from all patients before study entry. Animal experiments were approved by the Institutional Animal Care and Use Committee in Tsinghua University (18-CZJ2).

### Supplementary information

Supplementary data to this article can be found online at the appointed web site.

