## Supplementary figures and figure legends for "NOK promotes tumorigenesis through coordinating epidermal growth factor receptor to boost the downstream signaling in breast cancer"

**Supplementary figure legends:**

**Figure S1.** NOK expression was detected in its overexpression and knocked down cell lines. **A**, Over-expression of NOK in breast cancer cells by Ad-NOK. MDA-MB-231 cells were infected by an adenovirus expressing NOK (Ad-NOK). An adenovirus expressing GFP (Ad-GFP) was used as a control. A Western blot was performed to show the over-expression of NOK. GFP was used as a control for the equal transfection efficiency and b-actin was used as a loading control. **B**, NOK was stably depleted by two shRNAs in MDA-MB-231 cells. A Western blot analysis of endogenous NOK expression was performed for MDA-MB-231 cells stably infected by a lentivirus encoding two siRNAs against NOK. β-Actin was used as a loading control.

**Figure S2. NOK forms a complex with EGFR family members. A**, NOK-HA co-localized with EGFR. NOK-HA and EGFR-Flag were co-expressed in MDA-MB-231 cells. A microcopy analysis was performed to show the co-localization of NOK (red) and EGFR (green). An islet was enlarged to show the co-localization (yellow) of NOK and EGFR on the cell membranes. **B-C**, NOK associates with Her2 (B), Her3 (C) and Her4 (D). NOK-HA was co-expressed with Her2 (B), Her3 (C) and Her4 (D) in 293T cells. Cell lysates were immunoprecipitated with an antibody against Her2 (B), Her3 (C) or HA (D). The complexes were examined with indicated antibodies. The exogenously expressed proteins were examined in the cell lysates. The complex of immunoprecipitation was examined by Western blot analyses. **E**, The interaction of NOK with different mutations of EGFR is enhanced by EGF in 293 T cells. NOK and different EGFR mutates were over-expressed in 293T cells treated with or without EGF. Cell lysates were precipitated by an antibody against EGFR. The immunoprecipitated complexes were analyzed by a Western blot. **F**, The interaction of NOK with EGFR is not dependent on the phosphorylation of NOK. EGFR and different NOK mutants at tyrosine residues in the kinase domain were over-expressed in 293T cells. The immunoprecipitation analysis was performed using an antibody against Flag and the complexes were analyzed by a Western blot using an antibody against HA. The protein levels from different cells were showed in the lysates.

**Figure S3. NOK enhances EGF-induced colony formation and migration in breast cancer cells.** **A**, NOK enhances the soft agar colony formation in MDA-MB-231 induced by EGF. MDA-MB-231 cells were infected with Ad-NOK or Ad-GFP and seeded in 0.6% agar in 35 mm dishes with DMEM. Colonies were stained by crystal violet. **B**, Colonies >80 mm in diameter (Fig. S3A) were counted after 14 days. **C**, Over-expression of NOK promotes tumor cell migration. MDA-MB-231 cells were infected with Ad-NOK or Ad-GFP and seeded in transwell chambers with Matrigel. The migration cells were stained with crystal violet after 36 h culture in the presence or absence of EGF. **D**, Over-expression of NOK enhances migration cell numbers. The numbers of migration cells from the experiment in Fig. S3C were counted. * indicates the statistical significance. **E**, NOK mRNA was examined in wild type (WT) and NOK KO MEF cells. Different clones of cell lines were used. β-actin was used as a loading control. RT-PCR was performed using primers for the NOK and β-actin genes.

**Figure S4.** NOK enhances EGFR activation and the downstream signaling. **A-C**, NOK enhances the tyrosine phosphorylation of EGFR. NOK and EGFR were over-expressed in MDA-MB-231 cells (A), NIH-3T3 cells (B) and HEK293T cells (C) treated with or without EGF (100 ng/mL) for 5 min after starvation for overnight. Total tyrosine phosphorylated EGFR (pY-EGFR) was examined with a pY(4G10) antibody (C). Total EGFR levels were examined using an EGFR antibody (sc-03). **D**, The endogenous NOK was examined to show the efficiency of siRNAs against NOK in MDA-MB-231 cells transiently transfected with two siRNA oligos (NOKi, #1 and #2).

**Figure S5. The JM domain of NOK is critical for its activation on EGFR. A.** MDA-MB-231 cells were used to stably express NOK, NOK-JM and NOK-ΔJM. A vector was used as a control. Cells were treated with or without EGF. Phosphorylation of EGFR, STAT3, STAT5, and Erk1/2 was examined. Total EGFR, STAT3, STAT5 and β-actin were used as controls. The expression of NOK and its deletions was shown.

**TableS1.** A table of Pearson correlation of individual variants. Pearson correlations were calculated between NOK, EGFR, pSTAT5, pSTAT3, and Ki67,* indicated $p<0.05$ and ** indicated $p<0.01$.The total number of patients was 147.

**TableS2.A** table of Pearson correlation analysis of co-expression of NOK and EGFR top

STAT5 ,pSTAT3 and Ki67.Variants were defined as scores of co-expression of NOK and EGFR (NOK-EGFR), phosphorylation of STAT5 (pSTAT5) and STAT3 (pSTAT3), as well as Ki67.* indicated $p<0.05$ and ** indicated $p<0.01$ (n=147).

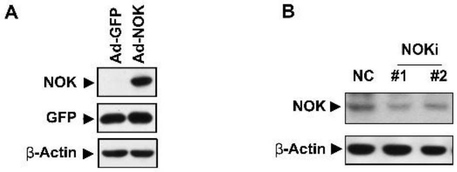


**Figure S1.**

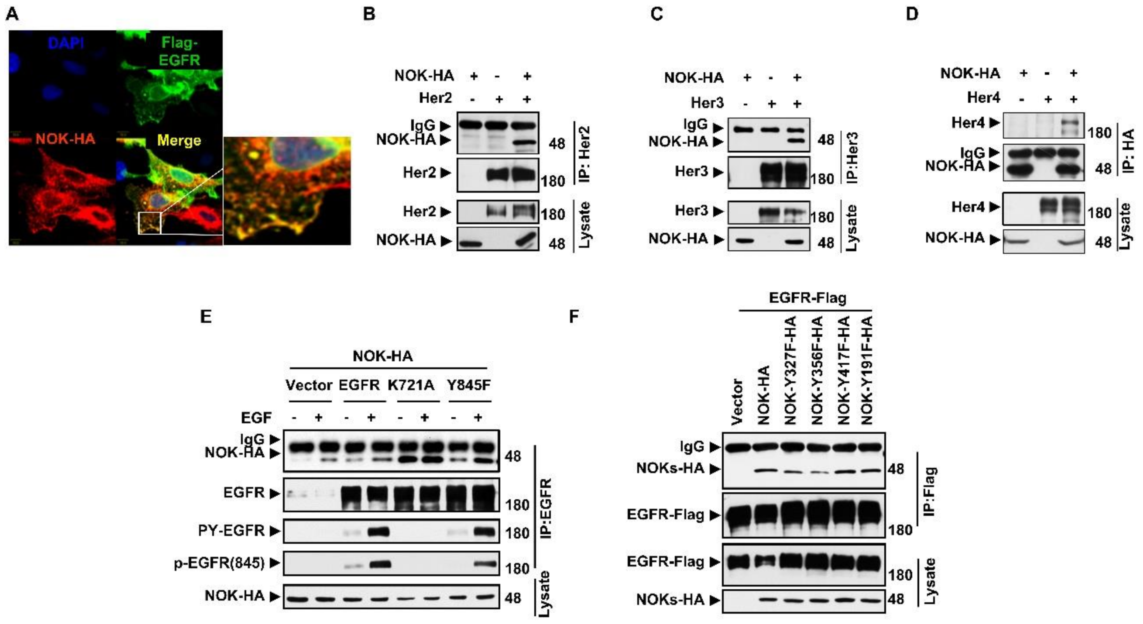
A
DAPI
Flag-EGFR
NOK-HA
Merge
B
NOK-HA + - +
Her2 - + +
IgG
NOK-HA
Her2
Her2
NOK-HA
48
180
180
48
IP: Her2
Lysate
C
NOK-HA + - +
Her3 - + +
IgG
NOK-HA
Her3
Her3
NOK-HA
48
180
180
48
IP:Her3
Lysate
D
NOK-HA + - +
Her4 - + +
Her4
IgG
NOK-HA
Her4
NOK-HA
180
48
180
48
IP: HA
Lysate
E
NOK-HA
Vector EGFR K721A Y845F
EGF - + - + - + - +
IgG
NOK-HA
EGFR
PY-EGFR
p-EGFR(845)
NOK-HA
48
180
180
180
48
IP:EGFR
Lysate
F
EGFR-Flag
Vector
NOK-HA
NOK-Y327F-HA
NOK-Y356F-HA
NOK-Y417F-HA
NOK-Y191F-HA
IgG
NOKs-HA
EGFR-Flag
EGFR-Flag
NOKs-HA
48
180
180
48
IP:Flag
Lysate

**Figure S2.**

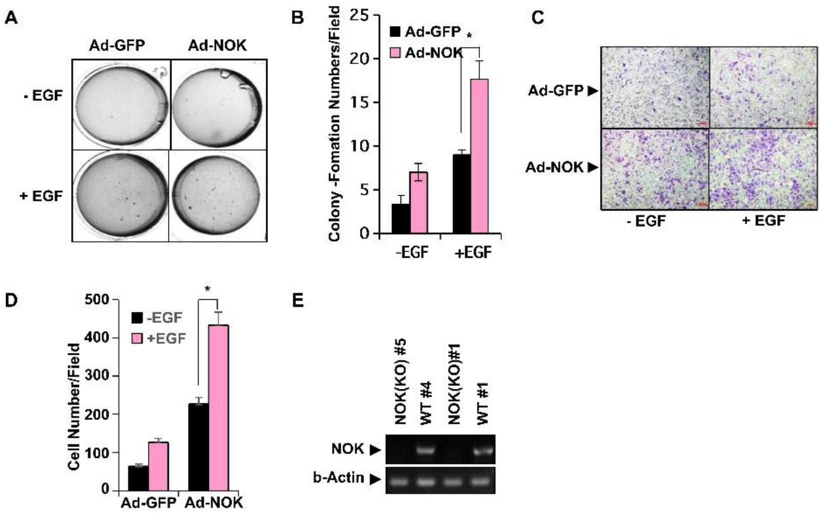
A
Ad-GFP
Ad-NOK
- EGF
+ EGF
B
Colony -Fomation Numbers/Field
Ad-GFP
Ad-NOK
*
0
5
10
15
20
25
-EGF
+EGF
C
Ad-GFP
Ad-NOK
- EGF
+ EGF
D
Cell Number/Field
-EGF
+EGF
*
0
100
200
300
400
500
Ad-GFP
Ad-NOK
E
NOK(KO) #5
WT #4
NOK(KO)#1
WT #1
NOK
b-Actin

**Figure S3.**

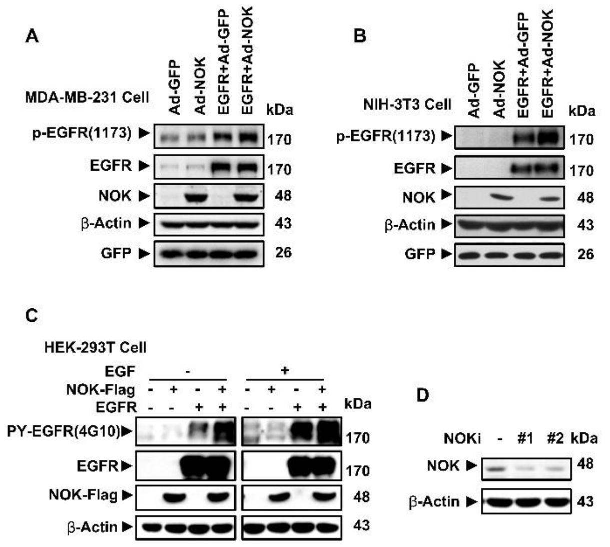
A
MDA-MB-231 Cell
Ad-GFP
Ad-NOK
EGFR+Ad-GFP
EGFR+Ad-NOK
kDa
p-EGFR(1173)
170
EGFR
170
NOK
48
β-Actin
43
GFP
26
B
NIH-3T3 Cell
Ad-GFP
Ad-NOK
EGFR+Ad-GFP
EGFR+Ad-NOK
kDa
p-EGFR(1173)
170
EGFR
170
NOK
48
β-Actin
43
GFP
26
C
HEK-293T Cell
EGF
-
+
NOK-Flag
EGFR
kDa
PY-EGFR(4G10)
170
EGFR
170
NOK-Flag
48
β-Actin
43
D
NOKi
-
#1
#2
kDa
NOK
48
β-Actin
43

**Figure S4.**

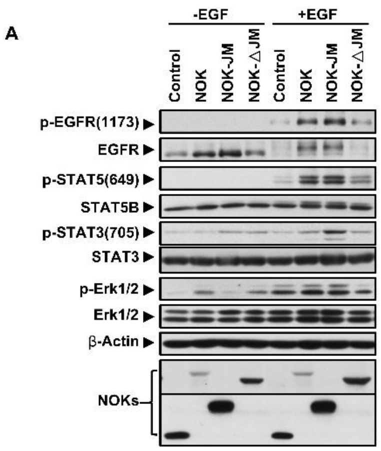
A
-EGF
+EGF
Control
NOK
NOK-JM
NOK-△JM
p-EGFR(1173)
EGFR
p-STAT5(649)
STAT5B
p-STAT3(705)
STAT3
p-Erk1/2
Erk1/2
β-Actin
NOKs

**Figure S5.**

### Suplementary table 1

**Pearson correlation analysis results**

| | | NOK_EGFR | pSTAT5 | pSTAT3 | Ki67 |
|---|---|---|---|---|---|
| **NOK_EGFR** | **Correlation** | **1** | 0.261** | 0.244** | 0.351** |
| | **Significance（Two tails）** | | 0.001 | 0.003 | 0 |
| | **N** | 147 | 147 | 147 | 147 |
| **pSTAT5** | **Correlation** | | **1** | 0.375** | 0.196* |
| | **Significance（Two tails）** | | | 0 | 0.017 |
| | **N** | | 147 | 147 | 147 |
| **pSTAT3** | **Correlation** | | | **1** | 0251* |
| | **Significance（Two tails）** | | | | 0.002 |
| | **N** | | | 147 | 147 |
| **Ki67** | **Correlation** | | | | **1** |
| | **Significance（Two tails）** | | | | |
| | **N** | | | | 147 |

**Supplementary table2**

**Pearson correlation analysis results**

| | | NOK | EGFR | pSTAT5 | pSTAT3 | Ki67 |
|---|---|---|---|---|---|---|
| **NOK** | **Correlation** | **1** | 0.117 | 0.349** | 0.316** | 0.439** |
| | **Significance （Two tails）** | | 0.158 | 0 | 0 | 0 |
| | **N** | 147 | 147 | 147 | 147 | 147 |
| **EGFR** | **Correlation** | | **1** | 0.133 | 0.138 | 0.285** |
| | **Significance （Two tails）** | | | 0.107 | 0.096 | 0 |
| | **N** | | 147 | 147 | 147 | 147 |
| **pSTAT5** | **Correlation** | | | **1** | 0.375** | 0.196* |
| | **Significance （Two tails）** | | | | 0 | 0.017 |
| | **N** | | | 147 | 147 | 147 |
| **pSTAT3** | **Correlation** | | | | **1** | 0.251** |
| | **Significance （Two tails）** | | | | | 0.002 |
| | **N** | | | | 147 | 147 |
| **Ki67** | **Correlation** | | | | | **1** |
| | **Significance （Two tails）** | | | | | |
| | **N** | | | | | 147 |
